# Mild traumatic brain injury/ concussion initiates an atypical astrocyte response caused by blood-brain barrier dysfunction

**DOI:** 10.1101/2021.05.28.446153

**Authors:** Benjamin P. Heithoff, Kijana K. George, Oleksii Shandra, Stefanie Robel

**Author notes:** Both authors contributed equally to the work (Benjamin P. Heithoff, Kijana K. George). **Corresponding Author:** Stefanie Robel, 2 Riverside Circle, Roanoke VA, 24016.

## Abstract

Mild traumatic brain injury/ concussion (mTBI) account for 70-90% of all reported TBI cases and cause long lasting neurological consequences in 10 to 40% of patients. Recent clinical studies revealed increased blood-brain barrier (BBB) permeability in mTBI patients, which correlated with secondary damage after mTBI. However, the cascade of cellular events initiated by exposure to blood-borne factors resulting in sustained damage are not fully resolved. We previously reported that astrocytes respond atypically to mTBI rapidly downregulating many proteins essential to their homeostatic function while classic scar formation does not occur. Here, we tested the hypothesis that mTBI -induced BBB damage causes atypical astrocytes through exposure to blood-borne factors. Using a mTBI mouse model, 2-photon imaging, an endothelial cell-specific genetic ablation approach, and serum-free primary astrocyte cultures, we demonstrated that areas with atypical astrocytes coincide with BBB damage and that exposure of astrocytes to plasma proteins is sufficient to initiate downregulation of astrocyte homeostatic proteins. While mTBI resulted in frequent impairment of both physical and metabolic BBB properties and leakage of small-sized blood-borne factors, deposition of the coagulation factor fibrinogen or vessel rupture were rare. Surprisingly, even months after mTBI BBB repair did not occur in areas with atypical astrocytes. Together, these findings implicate that even relatively small BBB disturbances are sustained long-term and render nearby astrocytes dysfunctional, likely at the cost of neuronal health and function.

## Introduction

Mild traumatic brain injury/ concussion (mTBI) make up 70-90% of all reported TBI cases, which are commonly seen in sports-related injury, car crashes, and blast exposure^1,2^. Of all patients who suffer from mTBI, 10-40% develop long-term neurological consequences that can significantly affect quality of life^3, 4^, such as cognitive impairment, depression, or sleep disturbances^5–7^.

Recent clinical studies have examined early biological events that may drive the progression of these neurological consequences after mTBI. With the use of dynamic contrast enhanced (DCE) magnetic resonance imaging (MRI), brain scans revealed increased bloodbrain barrier (BBB) permeability in patients that experienced a single mTBI^8, 9^. Increased BBB leakage correlates with secondary damage after mTBI, such as cerebral edema and increased hyperexcitability^10^, yet the characteristics of mTBI-induced BBB damage and subsequent cellular and molecular events leading to sustained damage after mTBI have yet to be elucidated.

Astrocytes respond to TBI by turning reactive and part of this response is shaped by exposure to blood-borne factors including the plasma proteins albumin^11–13^, thrombin^14, 15^, and fibrin^16^. Both thrombin and fibrin have been reported to trigger upregulation of glial fibrillary acidic protein (GFAP) in astrocytes, a classic indicator of astrogliosis. GFAP upregulation is part of the scar-forming response of astrocytes to focal injury, i.e. a site of primary tissue damage as a direct consequence of the TBI. Scar-formation protects uninjured brain areas of the brain from secondary damage induced by infiltration of immune cells or prolonged exposure to blood-borne factors^17, 18^.

Yet, we recently showed that mTBI/ concussion induced in mice by impact acceleration weight drop injury does not induce the classic scar-forming response of astrocytes characteristic for moderate and severe TBI, which have focal sites of injury surrounded by a glial scar. Instead, after mTBI, some astrocytes become mildly reactive but do not form scars and a separate subset of astrocytes respond atypically. Atypical astrocytes downregulate key astrocytic proteins, including glutamate transporter-1 (Glt1) and Kir4.1, within minutes after the injury in the absence of upregulation of GFAP. Despite the lack of most astrocyte-typical proteins, experiments using astrocyte-specific red fluorescent reporters demonstrated that the cells do not die^19^. However, the upstream mechanisms inducing the atypical astrocyte response after mTBI are not yet resolved.

Here we report that mTBI/concussion induced BBB damage in mice subjected to impact acceleration weight drop injury. Areas with BBB damage overlapped with atypical astrocytes. Thus, we hypothesized that mTBI/concussion-induced BBB damage causes atypical astrocytes through exposure to blood-borne factors. We assessed the extent of vascular and BBB damage using two-photon imaging and immunohistochemistry coupled with injection of tracer dyes. We deployed a genetic approach to mimic BBB damage in the absence of any other mechanical injury and astrocyte culture studies to test our hypothesis.

## Materials and Methods

### Mice

C57BL/6 mice of both sexes were used for weight drop injuries at 9 to 14 weeks of age. For endothelial cell-specific ablation, Gt(ROSA)26Sor^tm1(DTA)Jpmb^/J mice (Jackson Labs stock #006331) were crossed with Tg(Cdh5-cre/ERT2)1Rha (MGI:3848982). We refer to Gt(ROSA)26Sor^tm1(DTA)Jpmb^/J mice that express the Diphtheria Toxin A (DTA) subunit heterogeneously as DTA^(fl/wt)^ mice and Tg(Cdh5-cre/ERT2)1Rha that express the transgene heterozygously as Cdh5(PAC)-CreERT2^tg/wt^. To drive cre expression in adult mice, tamoxifen (10 mg/mL in corn oil) was administered to adult mice via oral gavage at 8.3 g/uL.

All animal procedures were approved and conducted according to the guidelines of the Institutional Animal Care and Use Committee of Virginia Polytechnic and State University and were done in compliance with the National Institute of Health’s Guide for the Care and Use of Laboratory Animals.

### Animal procedures

An impact acceleration weight drop TBI model was used to model mTBI without focal injury as previously described^19^. Mice were anesthetized with 3% isoflurane for 5 min and then placed on a foam pad after the analgesic buprenorphine (0.05-0.1 mg/kg) was administered subcutaneously. A flat steel disc was placed on the mouse head to diffuse the impact across the cranium. A 100 g weight guided by a plexiglass tube was dropped from 50 cm height. For the 10 and 30 minutes post injury (mpi) timepoints, a single injury was performed (1xTBI). For all other timepoints that occurred days post injury (dpi), impacts were repeated (3 total) and occurred at a 45 minute inter-injury interval (3xTBI). Shams underwent all procedures with exception of the weight drop.

To assess BBB leakage, Cadaverine conjugated to Alexa Fluor-555 was injected retro-orbitally (0.33 mg/ mouse in 100 μL saline). 30 min later, mice were transcardially perfused with Phospho-Buffered Saline (PBS) followed by 4% paraformaldehyde (PFA).

Adlh-1l1-eGFP//FVB/N mice were used for repeated *in vivo* two-photon imaging via a thinned-skull cranial window as described previously^20^. Single mTBIs from 60 cm height (identical to 50 cm in key outcome measures) or repeated TBIs from 50 cm height were performed^19^.

### Serum-free primary astrocyte cultures

Magnetic-activated cell sorting (MACS) was used to isolate astrocytes as detailed in Holt, et al. 2016^21^ and according to manufacturer’s directions (Miltenyl Biotec, 134-042-401). Microglia and oligodendrocytes were removed first by using cell type-specific microbeads. A second magnetic separation was performed using astrocyte cell-specific antibody-2 (ACSA-2) microbeads.

Primary astrocytes were maintained in 1 mL 5X B27 serum-free media, prepared as described in **Table 1**. Astrocytes were plated on laminated coverslips at 200,000 cells per well in 48-well plates, and incubated at 37°C with 5% CO_2_ for 14 days. Fresh media was added for 2 days, replaced on day 3, and subsequently half the volume was replaced with fresh media every 3-4 days.

Blood from adult C57B/6 male mice was withdrawn via cardiac puncture and centrifuged at 14,000 rpm for 5 min at 4°C to remove erythrocytes and other cells contained in whole blood. The supernatant, containing plasma, was diluted to 10% in 5X B27 serum-free media and added to the coverslips for 24h. Heat-denaturation was performed at 75°C for 5 min.

### Immunohisto- and cytochemistry

Brains were post-fixed overnight in 4% PFA and cut coronally at 50 μm thickness. Primary antibodies are listed in **Table 2**. Immunohistochemistry and -cytochemistry were performed according to standard protocols as previously described^19, 22^. For ZO-1 staining, antigen retrieval was performed using 100 mg pepsin in 10mM hydrochloric acid (HCl) for 20 minutes at 37°C. Slices were then washed twice in PBST (PBS with 150 uL L-1 Tween-20) and incubated in 3% H_2_O_2_ for 10 minutes. After three PBS washes, slices were incubated in primary antibody solution for a minimum 48 hours at 4°C.

Images of mouse brain slices or coverslips were taken using a Nikon A1R confocal microscope with 10x or 20x air objectives, and Apo 40x/1.30 and 60x/1.40 oil immersion objectives. Quantifications were done in confocal images using Image J in the cortical gray matter as previously detailed^22^.3 to 5 slices were imaged and analyzed, one image per slice.

### Statistics

Statistics were calculated and graphed using GraphPad Prism 8 (GraphPad Software). Quantifications of CD45+ cells fall under a Poisson distribution instead of the standard Gaussian distribution because the number of CD45+ cells is reported as discrete data. In order to normalize and stabilize the variance of the data, the square root of the total number of CD45+ cells per brain was calculated followed by a one-factor ANOVA to compare groups.

All other data were tested for Gaussian distribution using the Kolmogorov-Smirnov (KS) normality test. Statistical tests were chosen accordingly and are specified in the results section or figure legend. For all graphs, data points are plotted as individual slices or coverslips in scatterplots, and mean with standard error of the mean (SEM). Data points from the same mouse or culture were plotted as dots of the same color. Statistics were also run after averaging per mouse, and p values differed but with no overall change in significant group differences. Statistical significance is indicated with *p≤0.05, **p≤ 0.01, ***p≤ 0.001, **** p≤ 0.0001.

## Results

### 4.1 Single mTBI/ concussion causes BBB damage in areas of atypical astrocytes

To determine if BBB leakage occurred after single mTBI, we used the small fluorescently-labeled molecule Cadaverine (<1 kDa), which detects even small disturbances in BBB function. While shams had little to no leakage in cortex, we observed distinct Cadaverine leakage across the cortical gray matter as early as 10 minutes post injury (mpi) **(Fig 1a-c)**. Cadaverine leakage overlapped with atypical astrocyte areas, identified by lack of Glt-1 immunolabeling^19^ **(Fig 1a)**. Here, we use Glt-1 as a readout for atypical astrocytes, after we confirmed downregulation of Kir4.1, S100b, glutamine synthetase, and Aldh1l1-eGFP in the same astrocytes in areas with Cadaverine leakage (data not shown). Cadaverine leakage continued at 1dpi after single and repeated mTBI (**Fig 1b-c)**. Because Cadaverine was injected 30min before mice were perfused, this data indicates that BBB disturbances persist at 1dpi. To estimate the extent of BBB leakage after mTBI, we stained for the large blood plasma protein fibrinogen (~340 kDa), which deposits outside of vessels as part of the coagulation response that halts bleeding. Fibrinogen deposition occurred sparsely in areas of atypical astrocytes after mTBI (4 of 6 mice, 5 of 18 slices had a single fibrinogen+ area) **(Fig 1d)** whereas Cadaverine leakage was detected in nearly every slice and often in multiple areas.

**Figure 1.**
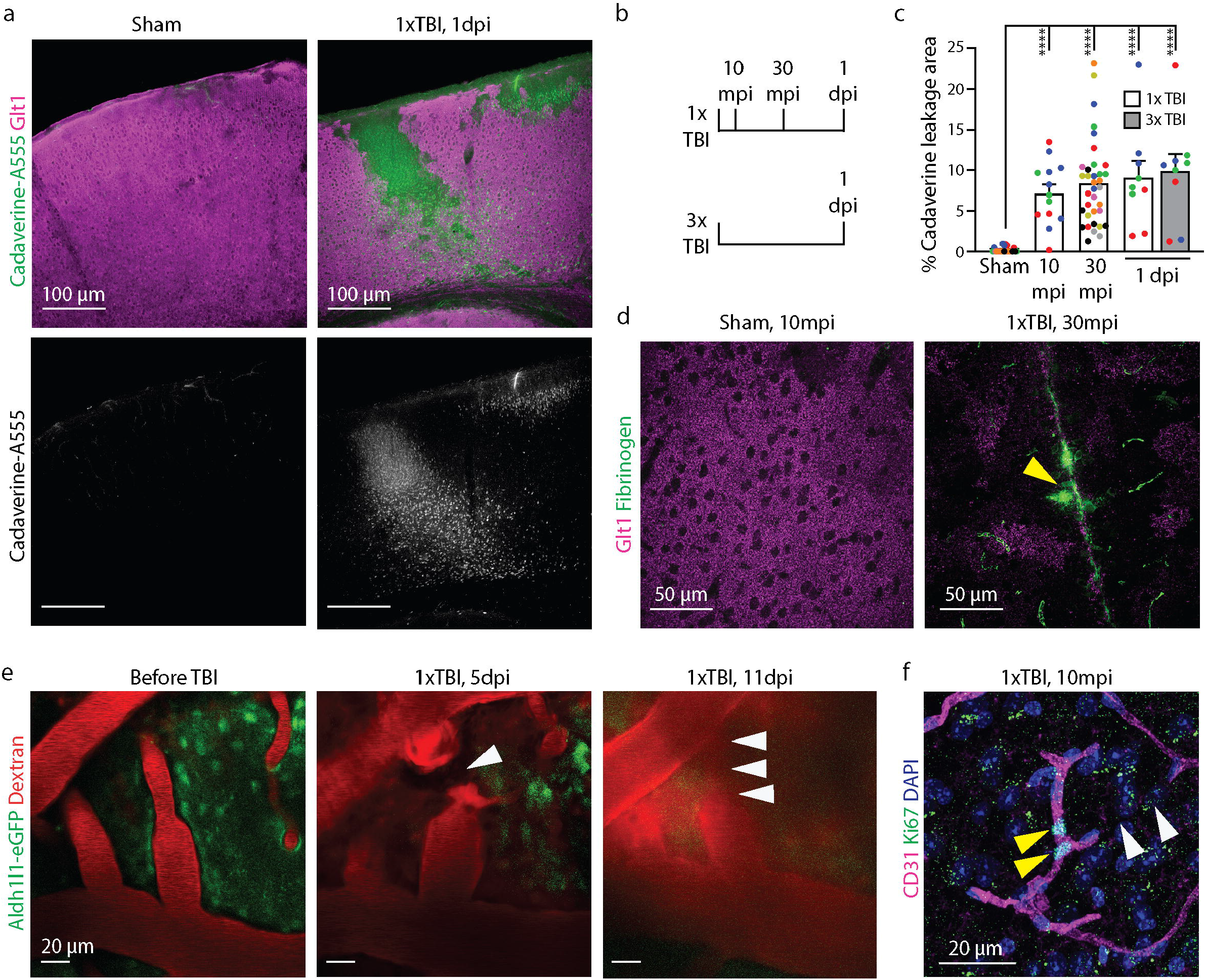
Blood-brain barrier leakage occurs in areas of atypical astrocytes after mild/ concussive TBI. **a.** Cadaverine leakage occurred in cortex by 10 minutes after a single mTBI and overlapped with regions of atypical astrocytes indicated by lack of expression of the glutamate transporter Glt1. **b.** Cadaverine was injected 30 minutes before perfusion for each timepoint. Single (1xTBI) and/or repeated (3xTBI) were induced at the timepoints listed. **c.** Cadaverine leakage was quantified as percent area of total cortex and plotted by slice. For 1dpi, both 1xTBI and 3xTBI were examined. Data are plotted by slice and are color-coded for each mouse. (Sham, 0.1963 ± 0.1506, n= 22; 1xTBI 10mpi, 7.168 ± 1.082, n= 13; 1xTBI 30mpi, 8.384 ± 0.8954, n= 35; 1xTBI 1dpi, 9.072 ± 2.086, n=9; 3xTBI 1dpi, 9.857 ± 2.126, n= 9. One-way ANOVA significant for post-injury timepoint, p< 0.0001; Dunn’s multiple comparisons test. Sham vs. 1xTBI 10mpi, p< 0.0001; Sham vs. 1xTBI 30mpi, p< 0.0001; Sham vs. 1xTBI 1dpi, p< 0.0001; Sham vs. 3xTBI 1dpi, p< 0.0001). **d.** Areas of fibrinogen deposition (yellow arrowhead), a large plasma protein, were found in sparse numbers in regions of atypical astrocytes indicated by lack of Glt1 expression. **e.** Live imaging of dextran-labeled vessels using two-photon microscopy revealed vessel rupture (white arrowheads) and reduced perfusion at 5 dpi. Reperfusion and repair of the same vessel (multiple white arrowheads) was observed at 11 dpi. **f.** The cell cycle marker Ki67 colocalized with the vessel marker CD31 (yellow arrowheads) in areas of abruptly short vessel patterns (white arrowheads), which indicated vessel proliferation. Of the 9 mice and 27 slices examined after mTBI, only 3 mice and 3 slices (one slice in each mouse) showed Ki67^+^/CD31^+^ colocalization. For 9 sham mice and 12 slices examined, no colocalization was detected. mpi = minutes post injury, dpi = days post injury.

Given that fibrinogen was detected in some slices, we next assessed direct damage to blood vessels. We used 2-photon imaging to image vessels before and after mTBI. In 1 of 10 regions of interest (ROIs) (n=3 mice), we observed vessel rupture acutely after mTBI, temporary loss of perfusion of the vessel at 5dpi, and subsequent reperfusion of the vessel at 11dpi (**Fig. 1e**). In 9 ROIs, we did not observe obvious vessel rupture after mTBI or leakage of the 70 kDa Dextran coupled to Tetramethylrhodamine (**Suppl. Fig. 1**). Because the area size that can be screened is limited to the imaging window in 2-photon imaging, we also used endothelial cell (EC) proliferation, which is required for repair of damaged vessels, as indirect readout for vessel damage across the cortical gray matter. As expected, very few Ki67+ cells were detected in the cortex of shams (data not shown), where only NG2 cells and microglia proliferate in the healthy adult brain. After mTBI, we occasionally found Ki67+/CD31+ EC (3/9 mTBI mice in 3/27 slices, 0/9 shams in 0/12 slices) close to vessels that appear disrupted, suggesting initiation of vessel repair (**Fig 1f)**.

Given that vessel rupture was observed rarely, we next asked if the BBB leakage after mTBI occurred due to impaired BBB properties. Zonula occludens-1 (ZO-1) is essential for forming tight junctions, which are responsible for maintaining a physically tight BBB. Vessels in sham mice had a continuous ZO-1 labeling pattern along CD31+ blood vessels while ZO-1 labeling along vessels near Cadaverine leakage was disrupted or absent at 1dpi (**Fig 2a-c)**. Another component of a functional BBB are transporters that ensure a transport of metabolites including glucose into the brain. To assess metabolic barrier properties, we examined the endothelial glucose transporter GLUT1 which accounts for 90% of brain glucose transport^23^. Vessels in shams were outlined by GLUT1 labeling, while GLUT 1 fluorescence intensity was decreased after mTBI **(Fig 2d,e)**. Taken together, mTBI resulted in frequent impairment of both physical and metabolic properties of the BBB and leakage of small-sized blood-borne molecules while larger vessel damage was rare.

**Figure 2.**
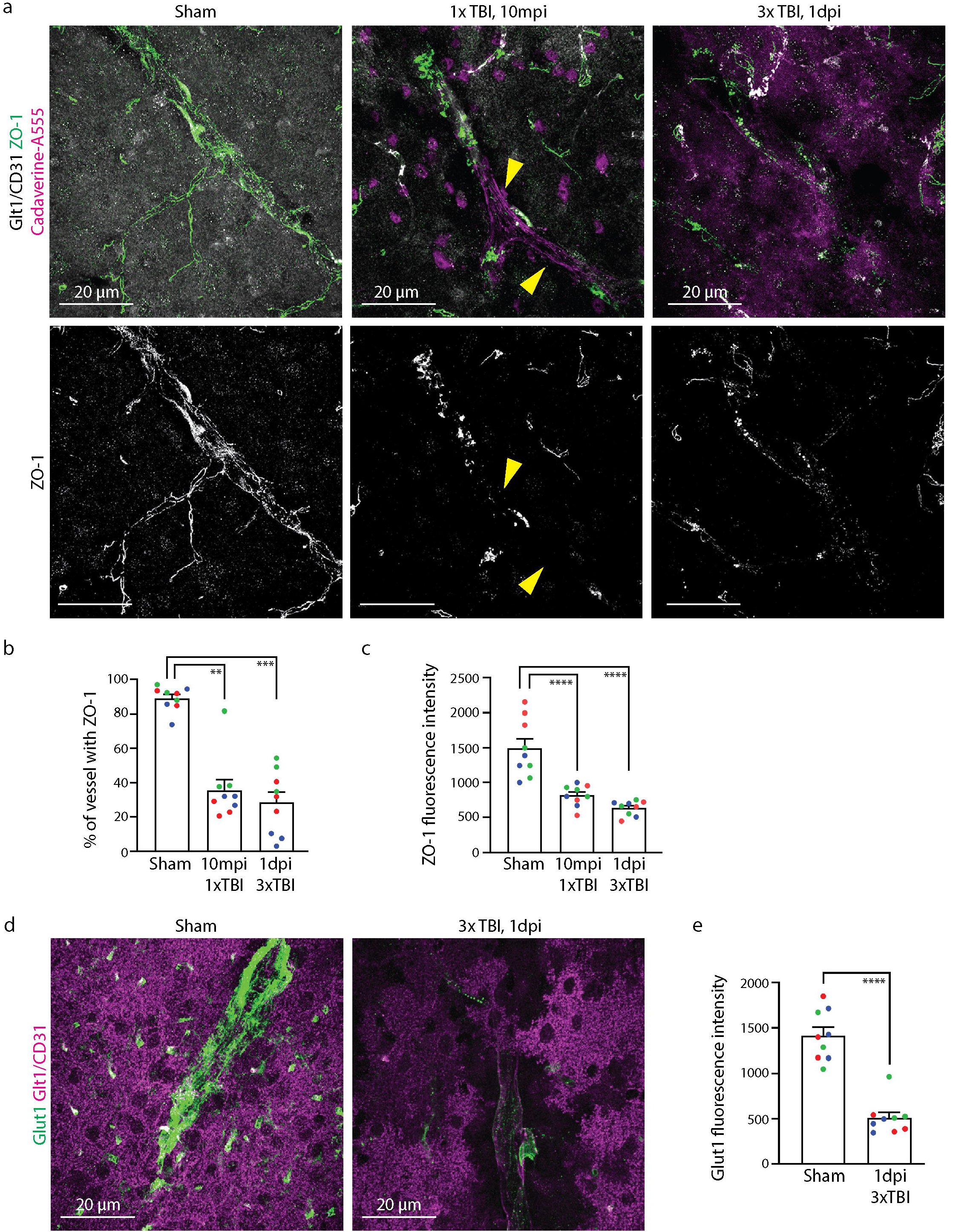
The blood-brain barrier is damaged after mTBI. **a.** Labeling of the tight junction protein ZO-1 was reduced and discontinuous after single and repeated mTBI, occurring in areas of atypical astrocytes (indicated by lack of Glt1 expression) and Cadaverine leakage. Some vessels completely lacked ZO-1 labeling (yellow arrowheads). **b.** Continuity in ZO-1 labeling of vessels was quantified by binarizing ZO-1 signal and drawing a line along the vessel using the vessel marker CD31 as a guide. The percentage of pixels with ZO-1 was calculated as percent of vessel with ZO-1 and was plotted by slice. Data are plotted by slice and are color-coded for each mouse. (Sham, 89.05 ± 2.329, n= 9; 1xTBI 10mpi, 35.65 ± 6.101, n= 9; 3xTBI 1dpi, 28.33 ± 6.150, n= 9. One-way ANOVA significant for post-injury timepoint, p= 0.0002. Dunn’s multiple comparisons test. Sham vs. 1xTBI 10mpi, p= 0.0019; Sham vs. 3xTBI 1dpi, p= 0.0006). **c.** Fluorescence intensity for the lines drawn in **b** were quantified and reported as mean grayscale (GS) units. (Sham, 1492 ± 137.7, n= 9; 1xTBI 10mpi, 814.7 ± 50.41, n= 9; 3xTBI 1dpi, 445.2 ± 35.54, n= 9; One-way ANOVA significant for post-injury timepoint, p< 0.0001. Tukey’s multiple comparisons test. Sham vs. 1xTBI 10mpi, p< 0.0001; Sham vs. 3xTBI 1 dpi, p< 0.0001). **d.** The endothelial glucose transporter GLUT1 was greatly reduced in its labeling at vessels after mTBI in areas of atypical astrocytes indicated by lack of expression of Glt1. **e.** Fluorescence intensity of GLUT1 was quantified after mTBI and reported as mean grayscale units. Data are plotted by slice and are color-coded for each mouse. (Sham, 1416 ± 92.46, n= 9; 3xTBI 1dpi, 508.8 ± 62.61, n= 9; Mann-Whitney Test. Sham vs. 3xTBI 1 dpi, p< 0.0001).

### 4.2 Exposure to blood plasma proteins is sufficient to induce atypical astrocytes in the absence of mTBI/ concussion

To determine if BBB leakage is sufficient to cause atypical astrocytes, we sparsely ablated ECs to open the BBB in the absence of other mechanical injury. Mice expressing the tamoxifen (Tx)-inducible CreERT2 recombinase driven by the EC-specific Cadherin 5 PAC promoter (Cdh5(PAC)) were crossed with mice expressing Diphtheria Toxin A (DTA) subunit behind a stop cassette flanked by loxP sites. A single low dose of Tx was administered to adult mice.

We observed EC ablation resulting in Cadaverine leakage within 2h post Tx administration (hpa) due to DTA-mediated suppression of protein synthesis^24^. As expected, we observed a significant decrease of ZO-1 covering CD31+ vessels after 2hpa, 6hpa, and 1dpa where endothelial cells were ablated. However, the fluorescence intensity of the remaining ZO-1 signal was unchanged when compared to controls hinting toward a different mechanism of tight junction loss in mTBI where overall ZO-1 intensity was reduced **(Fig 3a-c)**. GLUT1 expression was reduced at all timepoints **(Fig 3d,e)**.

**Figure 3.**
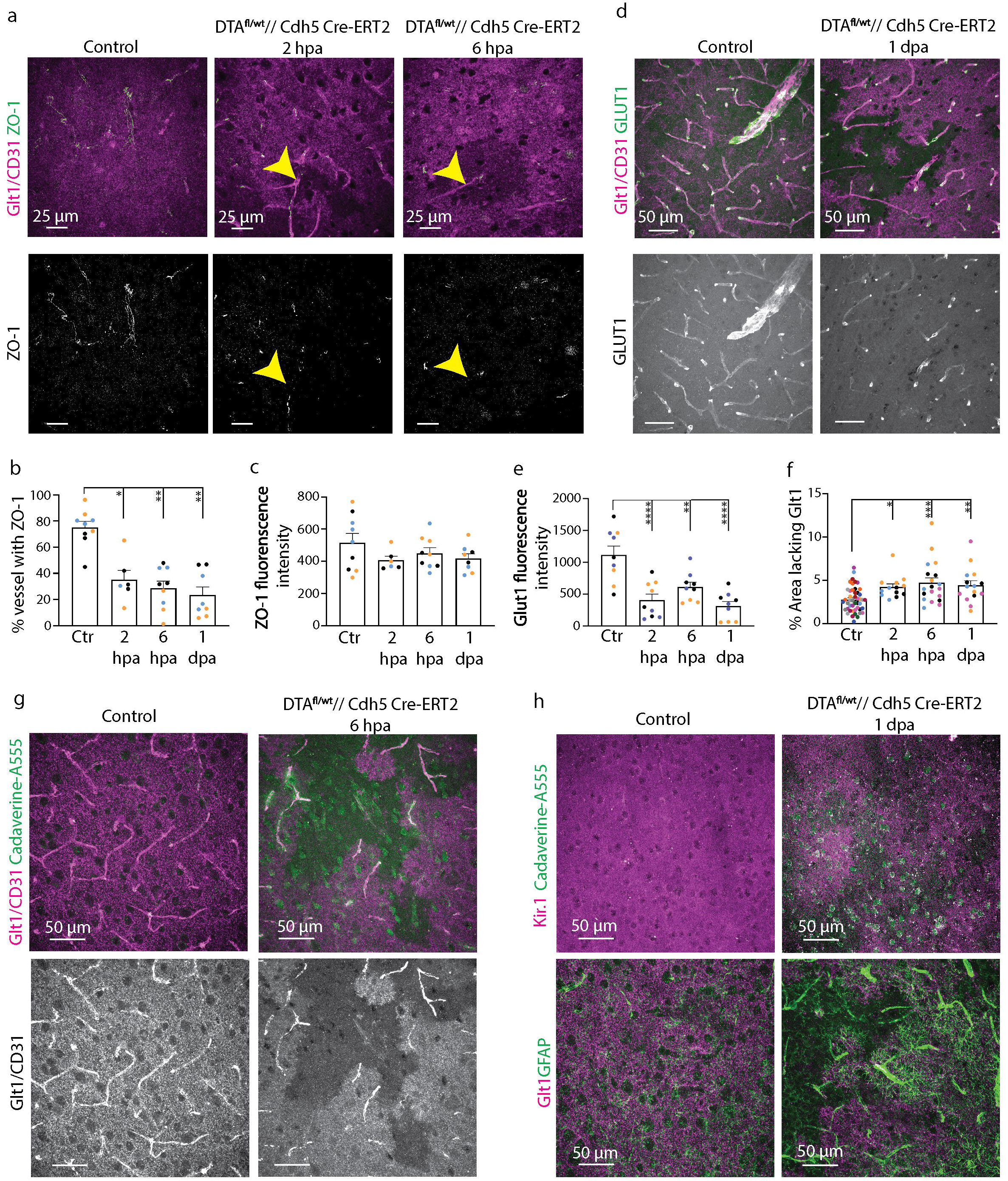
BBB leakage induced via endothelial-cell specific genetic ablation is sufficient to trigger atypical astrocytes in the absence of mTBI/ concussion. **a.** ZO-1 covering of CD31^+^ vessels was reduced as early as 2 hours after endothelial cell ablation and occurred in areas of atypical astrocytes, indicated by lack of Glt1 expression (yellow arrowheads). **b.** Continuity in ZO-1 labeling of vessels overlapping with atypical astrocytes was calculated as shown in **Figure 2b**. Data are plotted by slice and are color-coded for each mouse. (Control, 75.01 ± 4.745, n=9; 2hpa, 35.09 ± 7.089, n= 6; 6hpa, 28.71 ± 5.322, n= 9; 1dpa, 23.35 ± 6.198, n= 8. One-way ANOVA significant for post-administration timepoint. p< 0.0026. Dunnett’s multiple comparisons tests. p<0.0001. Control vs. 2hpa, p= 0.0386; Control vs. 6hpa, p=0.0024; Control vs. 1dpa, p=0.0029). **c.** Fluorescence intensity for the ZO-1 lines drawn in **b** were reported as grayscale values. Data are plotted by slice and are color-coded for each mouse. (Control, 517.1 ± 57.47, n= 9; 2hpa, 407.2 ± 25.39, n= 6; 6hpa, 451.9 ± 33.33, n= 9, 1 dpa, 418.1 ± 29.81, n= 9. Oneway ANOVA not significant for post-administration timepoint. p= 0.3907. Dunnett’s multiple comparisons test. p=0.2543. Control vs. 2hpa, p=0.2093; Control vs. 6hpa, p=0.5206; Control vs. 1dpa, p=0.2249). **d.** GLUT1 labeling was reduced at vessels in atypical astrocyte areas lacking Glt1 after genetically ablating endothelial cells after 2hpa**. e.** Fluorescence intensity of GLUT1 in areas of atypical astrocytes was calculated and reported as grayscale values. Data are plotted by slice and are color-coded for each mouse. (Control, 1117 ± 139.1, n= 9; 2hpa, 408.3 ± 93.00, n= 9; 6hpa, 612.1 ± 75.48, n= 9; 1dpa, 312.1 ± 72.25, n= 9. One-way ANOVA significant for post-administration timepoint. p= 0.0006. Dunnett’s multiple comparisons tests. p<0.0001. Control vs. 2hpa, p<0.0001; Control vs. 6hpa, p=0.0028; Control vs. 1dpa, p<0.0001). **f.** The percent area lacking Glt1 in cortex was quantified per slice at 2h, 6h, and 1d after endothelial cell ablation. Data points represent the percentage of area lacking Glt1 per slice in each group.(Control, 2.777 ± 0.091, n= 52; 2hpa, 4.234 ± 0.3619, n= 13; 6hpa, 4.716 ± 0.5636, n= 19; 1dpa, 4.417 ± 0.5277, n= 15. One-way ANOVA significant for postadministration timepoint. p= 0.004. Dunnett’s multiple comparisons tests. p=0.0002. Control vs. 2hpa, p=0.0271; Control vs. 6hpa, p=0.0004; Control vs. 1dpa, p=0.0072). **g.** A decrease in Glt1 labeling occurred as early as 2hpa and overlapped with areas of Cadaverine leakage **h.** Kir4.1 downregulation occurred in areas of atypical astrocytes (areas lacking Glt1), while GFAP upregulation occurred in areas adjacent to atypical astrocytes at 1 dpa. hpa= hours post administration, dpa= days post administration

In areas with Cadaverine leakage, astrocytes downregulated Glt1 and Kir4.1 consistent with our findings after mTBI 1dpa (**Fig 3f-h**). In contrast to our findings after mTBI, GFAP IHC was increased in astrocytes neighboring atypical areas at 1dpa (**Fig 3h**) suggesting that opening of the BBB is sufficient to induce atypical astrocytes^19^. To determine if blood-borne factors drive the changes in key homeostatic proteins in astrocytes after mTBI, we used a serum-free primary astrocyte culture model. Recent studies demonstrate that culturing astrocytes in chemically defined, serum-free medium enables astrocyte maturation while plasma-containing cultures remain immature^25^. Additionally, this assay allowed us to assess the effect of plasma on previously unexposed astrocytes. After maintaining primary astrocytes in serum-free media for 14 days, cultures were exposed to 10% fresh mouse plasma for 24h, which significantly reduced Glt-1 and Kir4.1 signal intensity. To determine if this was due to plasma protein or other plasma components (e.g. ions or lipids), plasma proteins were heat-denatured. Treatment of primary astrocytes with heat-denatured plasma rescued both Glt1 and Kir4.1 downregulation **(Fig 4a-c)** suggesting that plasma proteins initiate downregulation of astrocyte proteins in atypical astrocytes.

**Figure 4.**
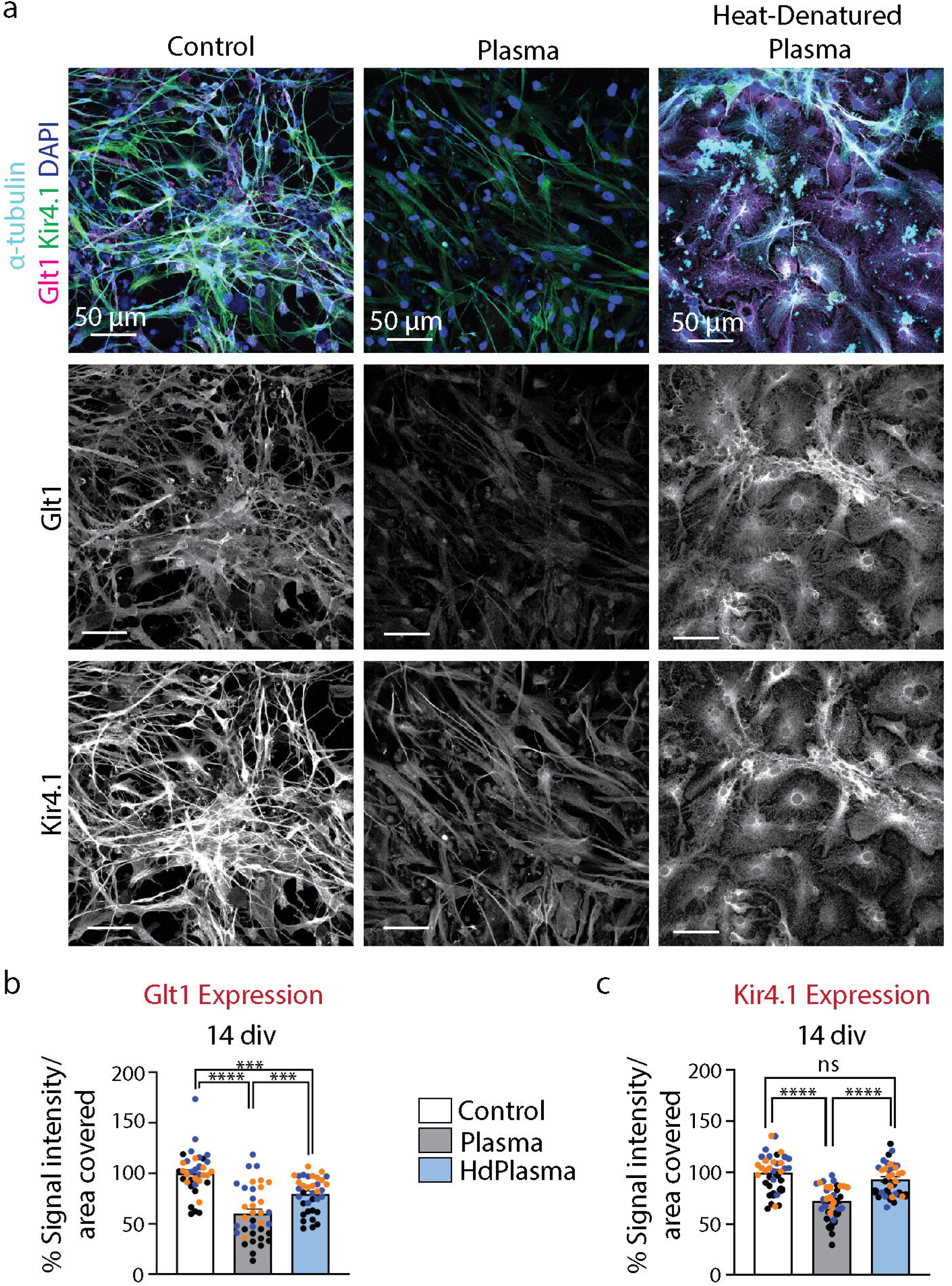
Glt1 and Kir4.1 expression in astrocytes is reduced 24h after plasma treatment *in vitro*. **a.** Treatment with heat-denatured plasma rescued both Glt1 and Kir4.1 expression in primary astrocyte cultures maintained in serum-free media for 14 *div*. **b,c.** Glt1 and Kir4.1 expression was quantified by dividing the mean gray value by the area coverage of each protein, then data were normalized in respect to the control group. Data points represent signal intensity for each ROI and are color-coded in respect to each independent culture. (*Glt1*. Control, 100 ± 3.603, n= 36 ROIs in 3 independent cultures; Plasma, 60.19 ± 4.429, n= 36 ROIs in 3 independent cultures; HdPlasma, 79.53 ± 2.85, n= 36 ROIs in 3 independent cultures. Oneway ANOVA significant for treatment. p< 0.0001. Tukey’s multiple comparisons test. p<0.0001. Control vs. Plasma, p<0.0001; Control vs. HdPlasma, p<0.0004; Plasma vs. HdPlasma, p=0.0010. *Kir4.1*. Control, 100 ± 3.033, n= 36 ROIs in 3 independent cultures; Plasma, 72.45 ± 2.651, n= 36 ROIs in 3 independent cultures; HdPlasma, 93.30 ± 2.62, n= 36 ROIs in 3 independent cultures. One-way ANOVA significant for treatment. p< 0.0001. Tukey’s multiple comparisons test. p<0.0001. Control vs. Plasma, p<0.0001. Control vs. HdPlasma, p=0.2071. Plasma vs. HdPlasma, p<0.0001). *div=* days in vitro, HdPlasma= Heat denatured plasma, ROI= region of interest

### 4.3 After mild TBI/ concussion the BBB does not repair in areas with atypical astrocytes

Previous studies demonstrate reduced entry of blood-borne factors within days after focal injury including stroke or moderate and severe TBI^26, 27^. In these studies, glial scars and/or repair of the BBB may contain BBB leakage. Scar formation does not occur after mTBI but the atypical astrocyte response is sustained for months after mTBI^19^. Thus, we asked if leakage of blood-borne factors is contained and whether the BBB is repaired in areas with atypical astrocytes.

Mice were injected with Cadaverine at 7dpi, 14dpi, and 64dpi. Cadaverine leakage persisted across all timepoints and leakage areas continued to overlap with atypical astrocytes **(5a-b)**. While single and repeated mTBI caused a similar extent of Cadaverine leakage at 1dpi, this area size was more than doubled in animals with repeated mTBI at 7dpi. This difference is largely due to a (not statistically significant) decrease of Cadaverine leakage from 1dpi to 7dpi **(Fig 5c)**. ZO-1 labeling was discontinuous and reduced in intensity at 64dpi (**Fig. 5f-h)** suggesting that sustained leakage may be due to lack of tight junction repair.

**Figure 5.**
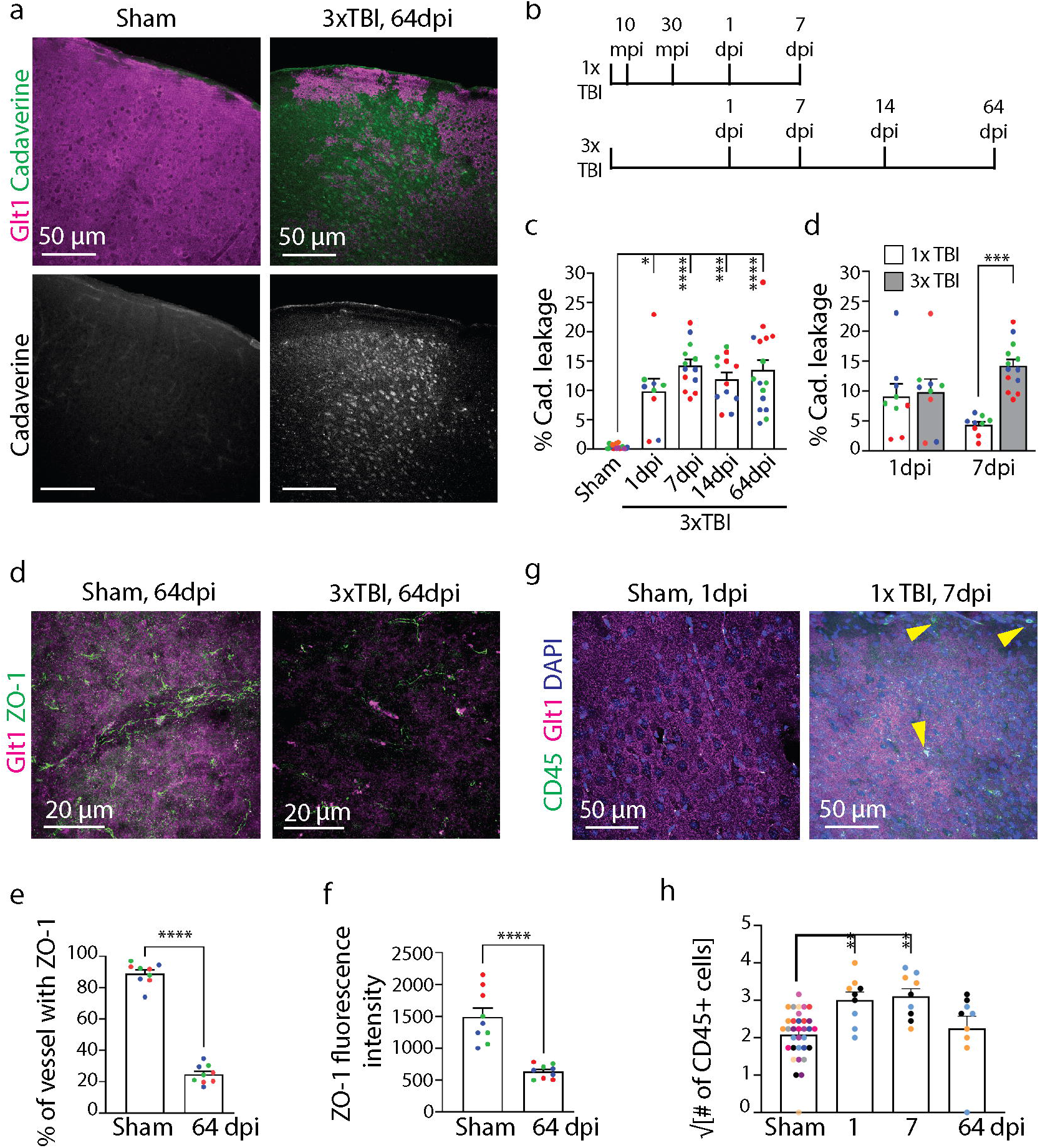
The BBB fails to repair in areas of atypical astrocytes. **a.** Cadaverine leakage was still present at 8 weeks after repeated mTBI and overlapped with areas of atypical astrocytes indicated by lack of Glt1 expression. **b.** Mice were examined at later timepoints after 3xTBI or 1xTBI (repeated, each injury separated by 45 min). Cadaverine was injected 30 minutes before sacrifice and perfusion at each timepoint. **c.** Cadaverine leakage was quantified at later timepoints after 3xTBI as percent area of total cortex and plotted by slice. Data are plotted by slice and are color-coded for each mouse. (Sham, 0.3444 ± 0.09558, n= 16; 3xTBI 1dpi, 8.955 ± 0.9901, n= 9; 3xTBI 7dpi, 14.18 ± 1.106, n= 13; 3xTBI 14dpi, 11.91 ± 1.183, n= 12; 3xTBI 64dpi, 13.48 ± 1.699, n= 16. Kruskal-Wallis test significant for post-injury timepoint, p< 0.0001. Dunn’s multiple comparisons test. Sham vs. 3xTBI 1dpi, p= 0.0112; Sham vs. 3xTBI 7dpi, p< 0.0001; Sham vs. 3xTBI 14dpi, p= 0.0002; Sham vs. 3xTBI 64dpi, p< 0.0001). **d.** Cadaverine leakage quantifications were compared between the 1dpi and 7dpi timepoints after either 1xTBI or 3xTBI. (1xTBI 1dpi, 8.955 ± 0.9901, n= 9; 1xTBI 7dpi, 4.315 ± 0.539, n= 9; 3xTBI 1dpi, 9.857 ± 2.126, n= 9; 3xTBI 7dpi, 3xTBI 7dpi, 14.18 ± 1.106. Two-way ANOVA significant for number of injuries, p= 0.0016, but not significant for post-injury timepoint, p= 0.8889; Tukey’s multiple comparisons test. 1xTBI 7dpi vs. 3xTBI 7dpi, p= 0.0002; 1xTBI 1dpi vs. 3xTBI 1dpi, p= 0.9859; 1xTBI 1dpi vs. 1xTBI 7dpi, p= 0.1802; 3xTBI 1dpi vs. 3xTBI 7dpi, p= 0.1893). **e.** Continuity of ZO-1 labeling along vessels was quantified at 64dpi 3xTBI in areas of atypical astrocytes as shown in **Fig 2b.** Data are plotted by slice and are color-coded for each mouse (Sham, 11.02 ± 2.602, n= 3; 3xTBI 8 wpi, 75.08 ± 1.531, n= 3. Unpaired t test for Sham vs. 3xTBI 8 wpi, p< 0.0001). **f.** Fluorescence intensity for the ZO-1 lines drawn in **e** and were reported as grayscale values. (Sham, 1491 ± 250.9, n= 3; 3xTBI 8 wpi, 522.4 ± 63.87, n= 3; Unpaired t test for Sham vs. 3xTBI 8 wpi, p= 0.0201). **g.** Clusters of CD45^+^ cells appeared in cortex after mTBI, while CD45^+^ in sham mice were found localized to meninges and not in cortex. **h.** Immune cell infiltration was quantified by taking the square root of the total number of CD45^+^ cells per brain slice. Data points represent the square root of the total number of CD45^+^ cells per slice and are color-coded for each mouse. (Sham, 2.083 ± 0.1141, n = 33, 3; 1dpi, 3.016 ± 0.2091, n= 9; 7 dpi, 3.112 ± 0.1989, n= 9; 64 dpi, 2.250 ± 0.3214, n= 9. One-way ANOVA significant for postinjury timepoint. p< 0.0001 Tukey’s multiple comparisons tests. p=0.0002. Sham vs. 1 dpi, p=0.0002; Sham vs. 7 dpi, p=0.0013, Sham vs. 64 dpi, p=0.92; 1 dpi vs. 7 dpi, p=0.9912; 1 dpi vs. 64 dpi, p=0.1032; 7 dpi vs. 64 dpi, p=0.053).

To determine if this prolonged BBB leakage resulted in immune cell infiltration, we used immunohistochemistry against CD45, which labels all lymphohematopoietic cells^28^. While CD45^+^ cells found in shams were restricted to the meninges, known to contain immune cells, we observed 5-10 cells within the cortical parenchyma at 10, 30mpi, and 7dpi after mTBI. Values were comparable to shams by 64 wpi (**Fig 5d-e**), indicating that BBB damage after mTBI/ concussion, in contrast to moderate and severe TBI, does not result in massive CD45 cell infiltration^29, 30^.

## Discussion

Our data demonstrate that even mTBI causes BBB damage characterized by leakage of small tracers and disrupted tight junctions in almost all mice that were examined. Fibrinogen, a large plasma protein (340 kDa) responsible for clotting and suggestive of bleeding after more severe vessel damage, was observed at much lower frequencies^31^. We also found only a mild increase of CD45+ immune cells after mTBI. This is in line with clinical data demonstrating that microbleeds occur only in a subset of mTBI patients. Microbleeds are small areas of hyperintensity in T2-weighted magnetic resonance imaging likely caused by iron-laden peripheral macrophages^32^. There is controversy around the association of microbleeds with worse mTBI outcomes as some studies report increased disability while others demonstrate no difference in post-concussion symptoms^33, 34^. We note significant differences in those studies with regard to the classification of mTBI (some studies include patients with focal injury based on high Glascow Coma Scale scores), outcome measures, and study size. However, an alternative explanation supported by our study might be that microbleeds represent only a small fraction of the actual vascular damage, which exposes the brain to many blood-borne factors including smaller size plasma proteins (albumin 67 kDa, thrombin 36 kDa) in the absence of hemorrhage or immune cell infiltration.

In the absence of glial scar formation, BBB leakage persisted unrestrained for 2 months after repeated mTBI. This is in contrast to focal TBI where leakage is contained after 10-14 days^26^. In our study, we demonstrate a lack of tight junction repair in areas with atypical astrocytes, yet it remains unresolved whether vessels damaged by brain injury have the innate ability to repair the BBB. Interestingly, evidence for tight junction re-formation is limited to cell culture studies in non-neural tissue^35^. There are no *in vivo* studies demonstrating re-appearance of tight junctions in previously damaged vessels. We recently demonstrated that astrocytes are necessary for maintenance of the BBB in the adult brain using a tamoxifen-inducible model of astrocyte ablation^22^. Because atypical astrocytes downregulate all astrocyte-specific proteins that we tested for^19^ (data not shown). Thus, an alternative interpretation to the EC-intrinsic failure to repair the BBB is that loss of function of atypical astrocytes results in lack of BBB repair.

Our endothelial cell ablation studies indicate that exposure of astrocytes to blood-borne factors is sufficient to induce the atypical response independently of mechanical strain induced by mTBI/ concussion. Because this effect is mimicked by exposure of astrocyte cultures to blood plasma and expression of astrocyte proteins is restored by heat-denaturation of the plasma, we suggest that plasma proteins initiate this unusual response. Previous studies using severe TBI models reveal that albumin^36^ and fibrin^16^ induce TGFb signaling in astrocytes. Albumin causes secretion of the pro-inflammatory molecules transforming growth factor beta (TGF-B) and Interleukin-1-beta (IL-1B) in astrocytes^11–13^. Fibrin triggers GFAP upregulation in astrocytes^16^ similarly to thrombin which also drives glial fibrillary acidic protein (GFAP) along with matrix metalloproteinase-9 (MMP9) upregulation^14, 15^. This is in contrast to the atypical astrocyte response after mild TBI/ concussion where GFAP is not expressed and other astrocyte proteins are downregulated. It is possible that the specific injury context, conditions within the extracellular environment, and combination of factors to which astrocytes are exposed dictate the response via specific signaling. In fact, the EC-ablation experiments hint toward this given that GFAP is upregulated in this context. Here, more severe vessel damage similar to vessel rupture is mimicked given that entire ECs are removed enabling entry of larger bloodborne factors. The specific signaling mechanism downstream of exposure to blood-born factors initiating the atypical astrocyte response remains to be elucidated in future studies.

## Conclusion

An increasing number of pre-clinical and basic research studies point to blood-brain barrier (BBB) damage as a risk factor for long-term neurological deficits after mTBI and for neurodegeneration. Here we conclude that mTBI/ concussion causes BBB damage that exposes astrocytes to plasma proteins initiating an atypical response characterized by severely reduced expression of astrocyte-typical homeostatic proteins. This is likely one cause of the lack of BBB repair after mTBI, thus playing important roles in the secondary injury cascade initiated by TBI.

## Acknowledgements

We thank Dr. Carmen Muñoz-Ballester for helpful and insightful discussions regarding the manuscript.

## Authorship Confirmation Statement

This work has not been published or submitted for publication in any journal.

## Authorship Contribution

B.P.H., K.K.G., and S.R. designed and planned the experiments. B.P.H. and K.K.G. conducted experiments, and collected and analyzed data. O.S. contributed to two-photon experiments, including imaging and data analysis. B.P.H, K.K.G., and S.R. wrote the manuscript with input from all authors.

---

**Supplementary Figure 1.**
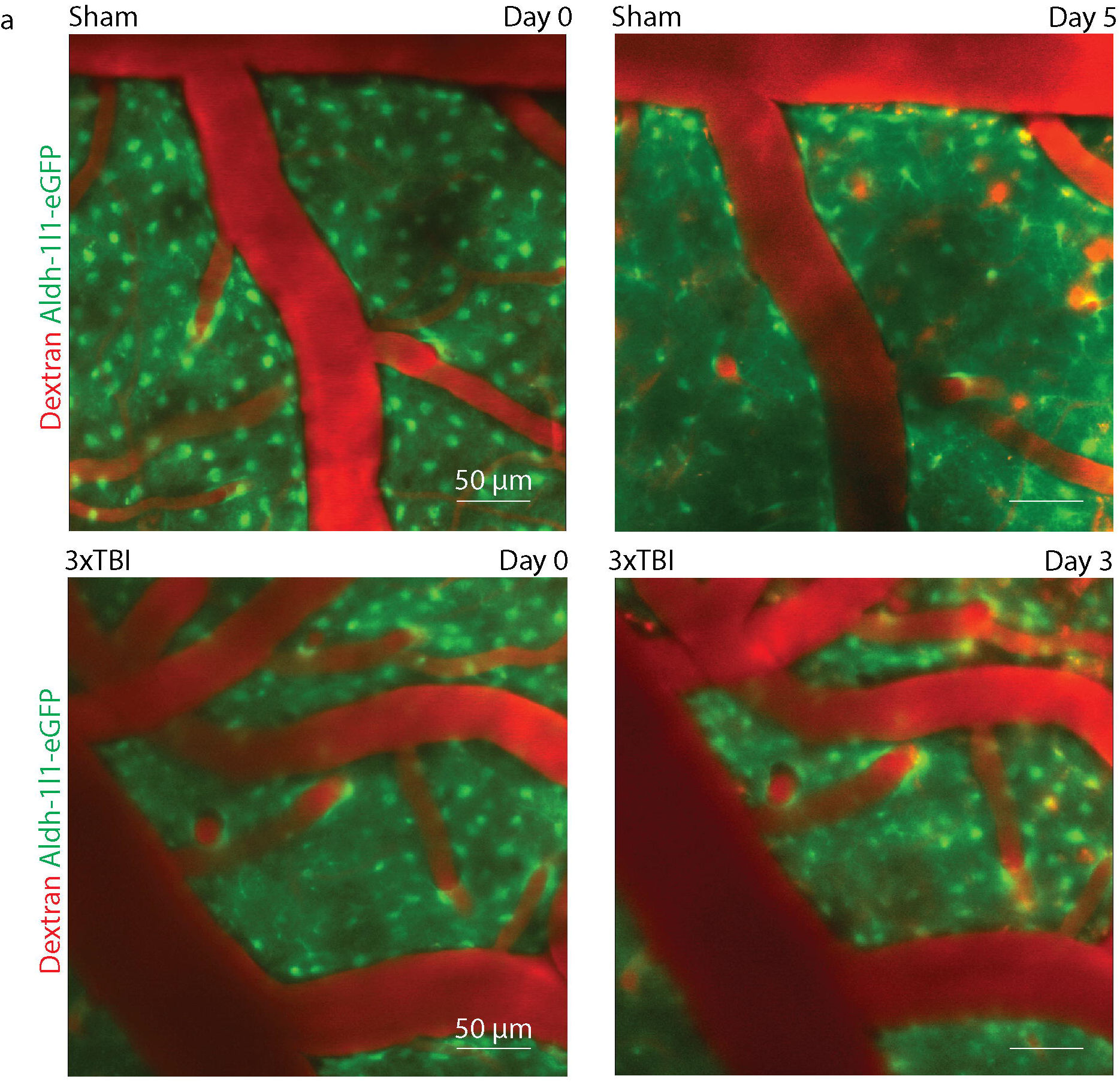
Intact blood vessels after mild TBI/ concussion in 9/10 ROIs. Live imaging of the cerebral vasculature after retro-orbital injection of a 70kDa Dextran coupled to Tetramethylrhodamine using two-photon microscopy revealing intact vasculature and perfusion at all timepoints (n=3, 9 ROI total, 3 mice were imaged before and acutely after repeated TBI, ¾ mice were subsequently imaged vessel). The figure depicts the final timepoint (3dpi). No leakage of the Dextran was observed in the Sham until day 5 (n=1, 3 ROIs).

## References

1. Cassidy, J.D., et al., Incidence, risk factors and prevention of mild traumatic brain injury: results of the WHO Collaborating Centre Task Force on Mild Traumatic Brain Injury. Journal of rehabilitation medicine, 2004. 36(0): p. 28–60.

2. Pavlovic, D., et al., Traumatic brain injury: neuropathological, neurocognitive and neurobehavioral sequelae. Pituitary, 2019. 22(3): p. 270–282.

3. Fehily, B. and M. Fitzgerald, Repeated Mild Traumatic Brain Injury: Potential Mechanisms of Damage. Cell transplantation, 2017. 26(7): p. 1131–1155.

4. Willer, B. and J.J. Leddy, Management of concussion and post-concussion syndrome. Curr Treat Options Neurol, 2006. 8(5): p. 415–26.

5. Bolton-Hall, A.N., W.B. Hubbard, and K.E. Saatman, Experimental designs for repeated mild traumatic brain injury: challenges and considerations. Journal of neurotrauma, 2019. 36(8): p. 1203–1221.

6. Ellis, M.J., et al., A Physiological Approach to Assessment and Rehabilitation of Acute Concussion in Collegiate and Professional Athletes. Frontiers in neurology, 2018. 9: p. 1115–1115.

7. Kenzie, E.S., et al., The Dynamics of Concussion: Mapping Pathophysiology, Persistence, and Recovery With Causal-Loop Diagramming. Frontiers in neurology, 2018. 9: p. 203–203.

8. Johnson, V.E., et al., Inflammation and white matter degeneration persist for years after a single traumatic brain injury. Brain, 2013. 136(Pt 1): p. 28–42.

9. Wang, M.L. and W.B. Li, Cognitive impairment after traumatic brain injury: The role of MRI and possible pathological basis. J Neurol Sci, 2016. 370: p. 244–250.

10. Sahyouni, R., et al., Effects of concussion on the blood-brain barrier in humans and rodents. Journal of concussion, 2017. 1: p. 10.1177/2059700216684518.

11. Ivens, S., et al., TGF-beta receptor-mediated albumin uptake into astrocytes is involved in neocortical epileptogenesis. Brain, 2007. 130(Pt 2): p. 535–47.

12. Cacheaux, L.P., et al., Transcriptome profiling reveals TGF-beta signaling involvement in epileptogenesis. J Neurosci, 2009. 29(28): p. 8927–35.

13. Hooper, C., et al., Differential effects of albumin on microglia and macrophages; implications for neurodegeneration following blood-brain barrier damage. J Neurochem, 2009. 109(3): p. 694–705.

14. Lin, C.C., et al., Thrombin mediates migration of rat brain astrocytes via PLC, Ca^2+^. CaMKII, PKCα, and AP-1-dependent matrix metalloproteinase-9 expression. Mol Neurobiol, 2013. 48(3): p. 616–30.

15. Mhatre, M., et al., Thrombin, a mediator of neurotoxicity and memory impairment. Neurobiol Aging, 2004. 25(6): p. 783–93.

16. Schachtrup, C., et al., Fibrinogen triggers astrocyte scar formation by promoting the availability of active TGF-beta after vascular damage. J Neurosci, 2010. 30(17): p. 5843–54.

17. Escartin, C., et al., Reactive astrocyte nomenclature, definitions, and future directions. Nat Neurosci, 2021. 24(3): p. 312–325.

18. Sofroniew, M. V., Molecular dissection of reactive astrogliosis and glial scar formation. Trends Neurosci, 2009. 32(12): p. 638–47.

19. Shandra, O., et al., Repetitive Diffuse Mild Traumatic Brain Injury Causes an Atypical Astrocyte Response and Spontaneous Recurrent Seizures. J Neurosci, 2019. 39(10): p. 1944–1963.

20. Shandra, O. and S. Robel, Imaging and Manipulating Astrocyte Function In Vivo in the Context of CNS Injury. Methods Mol Biol, 2019. 1938: p. 233–246.

21. Holt, L.M. and M.L. Olsen, Novel Applications of Magnetic Cell Sorting to Analyze CellType Specific Gene and Protein Expression in the Central Nervous System. PLoS One, 2016. 11(2): p. e0150290.

22. Heithoff, B.P., et al., Astrocytes are necessary for blood-brain barrier maintenance in the adult mouse brain. Glia, 2021. 69(2): p. 436–472.

23. Pardridge, W.M., R.J. Boado, and C.R. Farrell, Brain-type glucose transporter (GLUT-1) is selectively localized to the blood-brain barrier. Studies with quantitative western blotting and in situ hybridization. J Biol Chem, 1990. 265(29): p. 18035–40.

24. Ivanova, A., et al., In vivo genetic ablation by Cre-mediated expression of diphtheria toxin fragment A. Genesis, 2005. 43(3): p. 129–35.

25. Holt, L.M., et al., Astrocyte morphogenesis is dependent on BDNF signaling via astrocytic TrkB.T1. eLife, 2019. 8: p. e44667.

26. Bush, T.G., et al., Leukocyte Infiltration, Neuronal Degeneration, and Neurite Outgrowth after Ablation of Scar-Forming, Reactive Astrocytes in Adult Transgenic Mice. Neuron, 1999. 23(2): p. 297–308.

27. Simpkins, A.N., et al., Identification of Reversible Disruption of the Human Blood-Brain Barrier Following Acute Ischemia. Stroke, 2016. 47(9): p. 2405–2408.

28. Hendrickx, A. and X. Bossuyt, Quantification of the leukocyte common antigen (CD45) in mature B-cell malignancies. Cytometry, 2001. 46(6): p. 336–9.

29. Frik, J., et al., Cross-talk between monocyte invasion and astrocyte proliferation regulates scarring in brain injury. EMBO Rep, 2018. 19(5).

30. Kjell, J., et al., Defining the Adult Neural Stem Cell Niche Proteome Identifies Key Regulators of Adult Neurogenesis. Cell Stem Cell, 2020. 26(2): p. 277–293.e8.

31. Samuels, J.M., et al., Severe traumatic brain injury is associated with a unique coagulopathy phenotype. The journal of trauma and acute care surgery, 2019. 86(4): p. 686–693.

32. Griffin, A.D., et al., Traumatic microbleeds suggest vascular injury and predict disability in traumatic brain injury. Brain, 2019. 142(11): p. 3550–3564.

33. Huovinen, A., et al., Traumatic Microbleeds in Mild Traumatic Brain Injury Are Not Associated with Delayed Return to Work or Persisting Post-Concussion Symptoms. Journal of Neurotrauma, 2021.

34. Toth, L., et al., Traumatic brain injury-induced cerebral microbleeds in the elderly. GeroScience, 2021. 43(1): p. 125–136.

35. Chen, L., et al., Activating AMPK to Restore Tight Junction Assembly in Intestinal Epithelium and to Attenuate Experimental Colitis by Metformin. Front Pharmacol, 2018. 9: p. 761.

36. Senatorov, V.V., Jr., et al., Blood-brain barrier dysfunction in aging induces hyperactivation of TGFβ signaling and chronic yet reversible neural dysfunction. Sci Transl Med, 2019. 11(521).

