## Supplemental Tables 1-2 for "Mild traumatic brain injury/ concussion initiates an atypical astrocyte response caused by blood-brain barrier dysfunction"

**Table 1: Reagents to prepare 5x B27 serum-free media**

| **Serum Free Media** | |
| --- | --- |
| **To Make 500 mL of Media** | **Reagents** |
| 250 mL 1x NBM | 0.5x NBM |
| 250 mL 1x MEM | 0.5x MEM |
| 5 mL 100 mM sodium pyruvate | 1 mM sodium pyruvate |
| 5 mL 100 mM glutamine | 1 mM glutamine |
| 500 μL 5000U/mL pen/strep | 500 U pen/strep |
| 10 mL 50x B27 | 5x B27 |

Minimal Essential Media (catalog number: 51200038), NBM - Neurobasil Media (catalog number: 21103049),pen/strep - Penicillin-Streptomycin (catalog number: 15070063, 100x B27 (catalog number: 12587010), 100mMsodium pyruvate (catalog number:11360070), 200 mM glutamine (catalog number:25030081). All reagents were purchased from ThermoFisher Scientific.

**Table 2. List of antibodies**

| **Primary**  **Antibodies** |  |  |  |  |  |  |
| --- | --- | --- | --- | --- | --- | --- |
| **Name** | **Manufacturer** | **Catalog #** | **RRID** | **Species Raised in** | **Monoclonal/**  **Polyclonal** | **Concentration** |
| **S100β** | **Sigma Aldrich** | **S2532** | **AB_477499** | **Mouse** | **Monoclonal** | **1 : 1000** |
| **Glt1** | **Millipore** | **AB1783** | **AB_90949** | **Guinea pig** | **Polyclonal** | **1 : 1000** |
| **Ki67** | **Thermo Fisher** | **RM-9106-S1** | **AB_149792** | **Rabbit** | **Monoclonal** | **1 : 1000** |
| **Fibrinogen** | **Agilent** | **A008002** | **AB_578481** | **Rabbit** | **Polyclonal** | **1 : 500** |
| **ZO-1** | **Abcam** | **ab96587** | **AB-18.0006** | **Rabbit** | **Polyclonal** | **1 : 100** |
| **Iba1** | **Wako** | **#09-19741** | **AB_839504** | **Rabbit** | **Polyclonal** | **1 : 1000** |
| **Aquaporin-4 (AQ4)** | **Sigma Aldrich** | **A5971** | **AB_258270** | **Rabbit** | **Polyclonal** | **1 : 400** |
| **α-Tubulin** | **Sigma Aldrich** | **T6199** | **AB-19.0003** | **Mouse** | **Monoclonal** | **1:1000** |
| **CD45** | **Novus Biologicals** | **NB100-77417SS** | **AB-20.0005** | **Rat** | **Monoclonal** | **1:500** |
| **Beta-dystroglycan (β-dg)** | **Leica Biosystems** | **NCL-b-DG** | **AB_442043** | **Mouse** | **Monoclonal** | **1 : 200** |
| **Pan-laminin** | **Bio-Rad** | **AHP2491** | **AB_2133777** | **Rabbit** | **Polyclonal** | **1 : 1000** |
| **Collagen IV** | **Abcam** | **Ab6586** | **AB_305584** | **Rabbit** | **Polyclonal** | **1 : 1000** |
| **GLUT1** | **Novus Biologicals** | **NB300-666SS** | **AB_10000485** | **Rabbit** | **Polyclonal** | **1 : 1000** |

| **Secondary**  **Antibodies** |  |  |  |  |  |  |
| --- | --- | --- | --- | --- | --- | --- |
| **Name** | **Manufacturer** | **Catalog #** | **RRID** | **Species Raised in** | **Monoclonal/**  **Polyclonal** | **Concentration** |
| **Chicken Alexa-488** | **Jackson Immuno Research** | **703-546-155** | **AB_2340376** | **Donkey** | **Polyclonal** | **1 : 1000** |
| **Rabbit Alexa-488** | **Jackson Immuno Research** | **111-546-144** | **AB_2338057** | **Donkey** | **Polyclonal** | **1 : 1000** |
| **Mouse Alexa-488** | **Jackson Immuno Research** | **115-546-003** | **AB_2338859** | **Goat** | **Polyclonal** | **1 : 1000** |
| **Rat Alexa-488** | **Jackson Immuno Research** | **112-546-003** | **AB_2338364** | **Goat** | **Polyclonal** | **1 : 1000** |
| **Guinea pig Alexa-647** | **Jackson Immuno Research** | **106-606-003** | **AB_2337449** | **Goat** | **Polyclonal** | **1 : 1000** |
| **Rat Alexa-647** | **Jackson Immuno Research** | **712-605-153** | **AB_2340694** | **Donkey** | **Polyclonal** | **1 : 1000** |

| **Dyes** |  |  |  |  |  |  |
| --- | --- | --- | --- | --- | --- | --- |
| **Name** | **Manufacturer** | **Catalog #** | **RRID** | **Species Raised in** | **Monoclonal/**  **Polyclonal** | **Concentration** |
| **DAPI** | **ThermoFisher** | **D1306** | **AB_2629482** | **N /A** | **N /A** | **1 : 1000** |
| **Alexa-555 Cadaverine** | **ThermoFisher** | **A30677** | **N/A** | **N/A** | **N/A** | **0.33mg/mouse** |
| **Alexa-555 Dextran, 70,000 MW** | **ThermoFisher** | **D1818** | **N/A** | **N/A** | **N/A** | **0.33mg/mouse** |
